# The cation diffusion facilitator protein MamM’s cytoplasmic domain exhibits metal-type dependent binding modes and discriminates against Mn^2+^

**DOI:** 10.1101/2020.02.02.930644

**Authors:** Shiran Barber-Zucker, Jenny Hall, Afonso Froes, Sofiya Kolusheva, Fraser MacMillan, Raz Zarivach

**Affiliations:** Department of Life Sciences, Ben-Gurion University of the Negev, Beer Sheva 8410501, Israel; The National Institute for Biotechnology in the Negev, Ben-Gurion University of the Negev, Beer Sheva 8410501, Israel; Ilse Katz Institute for Nanoscale Science and Technology, Ben-Gurion University of the Negev, Beer Sheva 8410501, Israel; School of Chemistry, University of East Anglia, Norwich Research Park, Norwich NR4 7TJ, United Kingdom

**Author notes:** These authors contributed equally to the work (Shiran Barber-Zucker, Jenny Hall). Correspondence: Raz Zarivach; Fraser MacMillan.

**Keywords:** Cation diffusion facilitator, magnetotactic bacteria, metal selectivity, protein-metal interactions, electron paramagnetic resonance (EPR) spectroscopy, pulsed electron double resonance (PELDOR) spectroscopy

## Abstract

Cation diffusion facilitator (CDF) proteins are a conserved family of divalent transition metal cation transporters. CDF proteins are usually composed of two domains: the transmembrane domain (TMD), in which the metal cations are transported through, and a regulatory cytoplasmic C-terminal domain (CTD). Each CDF protein transports either one specific metal, or multiple metals, from the cytoplasm. Here, the model CDF protein MamM, from magnetotactic bacteria, was used to probe the role of the CTD in metal selectivity. Using a combination of biophysical and structural approaches, the binding of different metals to MamM CTD was characterized. Results reveal that different metals bind distinctively to MamM CTD in terms of; their binding sites, thermodynamics and binding-dependent conformation, both in crystal form and in solution. Furthermore, the results indicate that the CTD discriminates against Mn^2+^ and provides the first direct evidence that CDF CTD’s play a role in metal selectivity.

## Introduction

Cation diffusion facilitator (CDF) proteins are a conserved family of divalent transition metal cation transporters. CDF proteins form dimers, commonly homodimers, usually containing two domains: the N-terminal transmembrane domain (TMD) through which cations are transported, and the frequently found regulatory cytoplasmic C-terminal domain (CTD) (Coudray et al., 2013; Lopez-Redondo et al., 2018; Lu et al., 2009; Lu and Fu, 2007; Zeytuni et al., 2014b). Each CDF protein transports specific metal cations (either Zn^2+^, Mn^2+^, Cu^2+^, Co^2+^, Ni^2+^, Cd^2+^ or Fe^2+^) from the cytoplasm to either the extracellular environment, or inner-cellular compartments depending on its membranal location, typically by exploiting the proton-motive-force or by utilising other cation gradients (Barber-Zucker et al., 2017; Cubillas et al., 2013; Kolaj-Robin et al., 2015; Levy et al., 2019; Montanini et al., 2007). Since they control metal cation homeostasis at the cellular level, CDF proteins are crucial for proper cell function. As an example, malfunction of human CDF proteins leads to severe diseases including cardiovascular diseases, type-II diabetes, Parkinsonism and Alzheimer’s diseases (Huang and Tepaamorndech, 2013; Kambe et al., 2004). Although a greater understanding of CDF protein function at the molecular level has been achieved in recent years, the CDF transport mechanism is not yet fully understood. Furthermore, factors governing metal selectivity of these proteins are still under investigation (Barber-Zucker et al., 2017), hence the exact structure-function relationship of CDF proteins remains unresolved, and specifically how such relationships effect metal selectivity.

Magnetotactic bacteria (MTB) are a group of Gram-negative bacteria that align with and move along magnetic fields. The bacterium contains subcellular organelles named magnetosomes, each of which is composed of ~30-120 nm iron-based magnetic particle (magnetite or gregite) enclosed in a protein-rich lipid membrane (Barber-Zucker et al., 2016a; Komeili, 2012; Uebe and Schüler, 2016). The magnetosomes are arranged on an actin-like polymer in a chain arrangement, creating a magnetic dipole moment which allows these bacteria to use external magnetic fields. In nature, this property enables the bacteria to orient themselves to the geomagnetic field, thus navigating to their preferred habitat – usually the oxic-anoxic zones in aquatic environments (Barber-Zucker and Zarivach, 2017; Komeili et al., 2006). MamM, a CDF protein found uniquely on the magnetosome membrane, transports iron from the cytoplasm into the magnetosomes, enabling the synthesis of magnetic particles. It has been shown that deletion of the *mamM* gene, or even of its CTD, abolishes magnetic particle formation, emphasising its importance for the biomineralisation process (Uebe et al., 2011). In previous studies, a MamM CTD crystal structure was presented which adopts the characteristic fold of CDF protein CTDs, and the presence of which is crucial for the overall protein function. Additionally, three MamM CTD metal binding sites were proposed that included two symmetrical, allosteric peripheral sites (PSs, composed of H264 from one monomer and E289 from the second monomer) and one central site (CS, composed of D249 and H285 from both monomers). Biophysical evidence revealed that zinc binding to the PSs leads to a change in protein conformation from an open, dynamic V-shaped dimer to a much tighter, rigid dimeric packing (Barber-Zucker et al., 2016b; Barber-Zucker et al., 2019; Zeytuni et al., 2014b, 2014a). These studies and those of other bacterial CDF proteins suggest the following common mechanism for CDF proteins: at a certain cytoplasmic metal cation concentration, cations bind to the CTD, locking it in a tighter packing when compared to its apo V-shape. This, in turn, facilitates the conformational change of the TMD thus enabling cations to be transported through this domain to the other side of the membrane (Barber-Zucker et al., 2019; Cherezov et al., 2008; Cotrim et al., 2019; Kolaj-Robin et al., 2015; Lopez-Redondo et al., 2018; Zeytuni et al., 2014b). Although previous MamM studies characterised some of the roles and the mechanism of the CTD in CDF proteins, one question remained unanswered – whether the CTD has a role in metal selectivity.

In this study the well-characterised MamM CTD is used as a model to investigate the role of CTDs in metal selectivity *in vitro*, and this work presents, for the first time, structural evidence of metal binding of any CDF CTD to multiple metals. The crystal structure of MamM CTD was solved in the presence of Cu^2+^, Cd^2+^ and Ni^2+^, revealing the propensity of difference metals to bind at different metal-specific sites, leading to different conformations in the crystalline state. To further characterise the binding abilities of additional metal cations to MamM CTD in solution, tryptophan-fluorescence spectrometry, isothermal titration calorimetry (ITC) and pulsed electron-electron double resonance (PELDOR) spectroscopy were employed. Results indicate that MamM CTD can bind all the metal cations tested with the exception of Mn^2+^. Their binding commonly leads to the same or a similar tighter conformation, as that detected for the Cu^2+^-bound crystallographic conformation. Each bound metal, however, does exhibit different thermodynamic binding parameters implying a degree of selectivity for the effectivity of metal binding to the CTD. Overall, the results presented here suggest that MamM CTD discriminates against Mn^2+^, presumably to avoid impure iron-based magnetic particles synthesis. Thus, we see here the first direct evidence for the role of the CTD in metal selectivity in CDF proteins.

## Results and Discussion

### Crystal structures of MamM CTD with different metals exhibit different conformations and binding sites

The crystal structure of apo MamM CTD has been previously solved in two space groups, both revealing the same monomeric fold and dimerisation interface (pdb codes: 3W5X & 3W5Y) (Figure 1A-C, see RMSDs in Table 1 and distances and dihedral angels in Table 2) (Zeytuni et al., 2014b). To better achieve an understanding of the conformational changes occurring during metal binding in MamM (Barber-Zucker et al., 2019; Zeytuni et al., 2014b), co-crystallisation trials of MamM CTD (residues 215-318) with various divalent transition metal cations: Fe^2+^, Mn^2+^, Zn^2+^, Ni^2+^, Co^2+^, Cu^2+^ and Cd^2+^, under a range of different conditions were performed. Crystal structure models of MamM CTD were only obtained in the presence of three different metal cations: Cd^2+^, Ni^2+^ and Cu^2+^ (whilst crystals containing Zn^2+^ diffracted well, Zn^2+^ could not be detected in the electron density map). The structures with bound Cd^2+^ and Ni^2+^ indicate no conformational change both in their monomeric fold and their dimeric interface, as compared to the apo form (Figure 1A, C, Table 1 and Table 2). In contrast, the Cu^2+^-bound form reveals a clear change in the dimeric conformation to a tighter dimer interface, as has already been demonstrated in other CDF proteins and proposed for Zinc-bound MamM CTD in solution using PELDOR spectroscopy (Figure 1A, B, Table 1 and Table 2) (Barber-Zucker et al., 2019; Cherezov et al., 2008). Furthermore, in this specific Cu^2+^-bound structure, most of the C-terminal tail of one of the monomers, which was not previously identified in the apo forms’ electron density maps, is now resolved. The C-terminal residues (292-314) form a short β-strand which adds to the previously-solved three-stranded β-sheet, resulting in a 4-stranded β-sheet, followed by an α-helix. The C-terminal helix is found in the space between the two monomers and forms a stabilising bonding network with the adjacent monomer’s residues. This non-symmetric interaction may be due to crystal packing, or some as yet unknown biological function. The α-helical tail stabilises the CTD closed state and may interact with the TMD loops to also stabilise the transporter’s transmembrane conformation in its activated state. However, as CDFs homodimers have not previously revealed non-symmetric characteristics, we propose that these C-terminal interactions may be locking the dimer in its closed state, resulting in the stabilised crystal packing.

**Table 1:** RMSDs between different CDFs’ CTD dimeric structures.

|  | MamM apo<br>(3W5X) | MamM apo<br>(3W5Y) | MamM<br>Cu<sup>2+</sup>-bound<br>(6GP6) | MamM<br>Cd<sup>2+</sup>-bound<br>(6GMT) |
| --- | --- | --- | --- | --- |
| MamM apo (3W5X) | - | 0.82 | 1.56 | 0.35 |
| MamM apo (3W5Y) | - | - | 1.60 | 0.89 |
| MamM Cu<sup>2+</sup>-bound (6GP6) | - | - | - | 1.62 |
| MamM Cd<sup>2+</sup>-bound (6GMT) | - | - | - | - |
| MamM Ni<sup>2+</sup>-bound (6GMV) | 0.18 | 0.91 | 1.58 | 0.44 |
| CzrB apo (3BYP) | 1.49 | 1.59 | - | - |
| CzrB bound (3BYR) | - | - | 1.43 | 1.64 |
| EcYiiP bound (3H90) | - | - | 1.33 | 1.19 |
| SoYiiP bound (5VRF)<sup>a</sup> | - | - | - | - |
| MamB apo (5HO5) | 1.21 | 1.20 | 1.32 | 1.26<sup>b</sup> |
| MamB bound (5HO1) | 1.76 | 1.62 | 1.37 | 1.13 |

All RMSDs between the dimeric structures were calculated using the iterative magic fit tool of Swiss-PdbViewer 4.1.0 (Guex and Peitsch, 1997).
<sup>a</sup> Satisfying fit directly between the dimeric MamM and SoYiiP structures could not be obtained.
<sup>b</sup> RMSD was calculated using the domain fragment alternate fit tool of Swiss-PdbViewer 4.1.0.

**Table 2:** Dihedral angles and distances between selected residues in all CDFs’ CTD structures.

|  | Distance (Å) |  |  | Dihedral angle (°) |  |
| --- | --- | --- | --- | --- | --- |
|  | R240-R240 | G276-G276 | V242-V242 | R240-P256-P256-R240 | V242-V260-V260-V242 |
| MamM apo<br>(3W5X) | 25.78 | 45.40 | 30.03 | 48.759 | -47.116 |
| MamM apo<br>(3W5Y) | 22.86 | 44.31 | 27.34 | 43.240 | -41.640 |
| MamM Cu<sup>2+</sup>-<br>bound (6GP6) | 21.47 | 39.57 | 19.39 | 28.533 | -34.863 |
| MamM Cd<sup>2+</sup>-<br>bound<br>(6GMT) | 26.18 | 45.11 | 29.23 | 48.110 | -48.105 |
| MamM Ni<sup>2+</sup>-<br>bound<br>(6GMV) | 26.34 | 45.51 | 30.01 | 50.176 | -47.742 |
|  | R234-R234 | Q270-Q270 | A236-A236 | R234-G250-G250-R234 | A236-V254-V254-A236 |
| CzrB apo<br>(3BYP) | 28.13 | 45.54 | 30.96 | 51.849 | -51.592 |
| CzrB bound<br>(3BYR) | 12.22 | 37.60 | 13.16 | 21.354 | -14.796 |
|  | R237-R237 | R273-R273 | S239-S239 | R237-D253-D253-R237 | S239-L257-L257-S239 |
| EcYiiP bound<br>(3H90) | 14.26 | 36.86 | 13.89 | 23.880 | -16.838 |
|  | R239-R239 | A275-A275 | A241-A241 | R239-G255-G255-R239 | A241-L259-L259-A241 |
| SoYiiP bound<br>(5VRF) | 15.28 | 39.11 | 15.19 | 23.801 | -17.963 |
|  | R238-R238 | A274-A274 | V240-V240 | R238-A254-A254-R238 | V240-V258-V258-V240 |
| MamB apo<br>(5HO5) | 19.44 | 39.79 | 17.10 | 28.135 | -30.799 |
| MamB bound<br>(5HO1) | 19.10 | 38.93 | 18.25 | 26.612 | -32.822 |

All distances and dihedral angles were measured between the residues' $C_{\alpha}$ from both monomers, using UCSF Chimera package, version 1.12 (Pettersen et al., 2004).
All the residues in each column are found in the same location in all the proteins.

**Figure 1:**
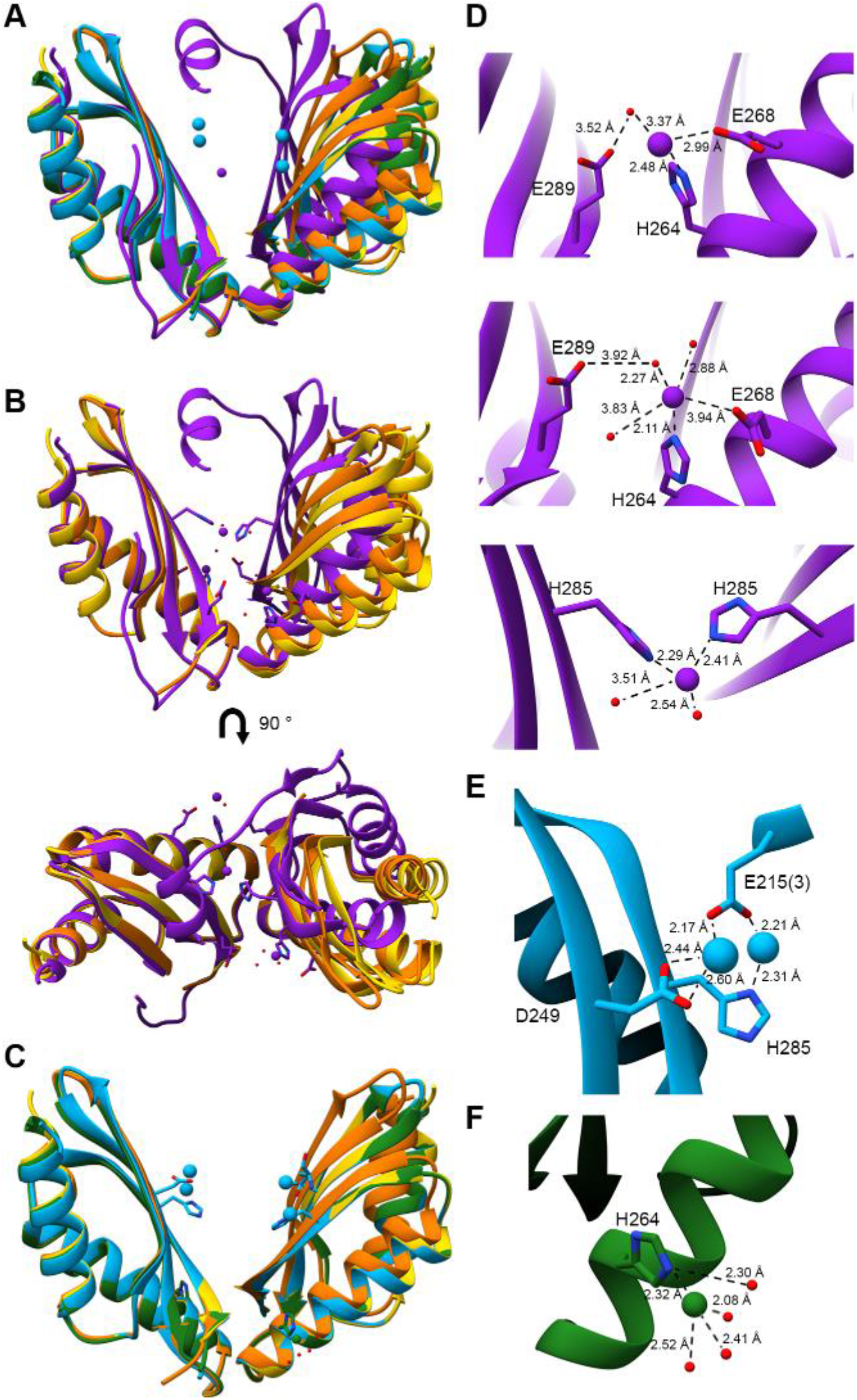
Crystal structures of MamM CTD with different metal cations. (A) The three MamM CTD bound structures (Cd^2+^ (pdb code: 6GMT) – blue, Ni^2+^ (pdb code: 6GMV) – green, Cu^2+^ (pdb code: 6GP6) – purple) and the two apo structures (pdb code: 3W5X in yellow, pdb code: 3W5Y in orange (Zeytuni et al., 2014b)), overlapped onto each other. Metal cations are presented for each bound structure in the same colours. (B) Apo forms and Cu^2+^-bound structures overlapped onto each other from two angles, showing the conformational change exhibited by the Cu^2+^-bound structure. The metal binding residues are presented as sticks. (C) Apo forms, Ni^2+^-bound and Cd^2+^-bound structures overlapped onto each other, indicating no conformational changes between all structures. The metal binding residues are presented as sticks. (D) Close-up of the Cu^2+^-bound structure binding sites (upper and middle panels- PSs, lower panel-CS). (E) Close-up of the Cd^2+^-bound structure binding site (of one monomer). (3) refers to a non-biological- assembly monomer. (F) Close-up of the Ni^2+^-bound structure binding site (of one monomer). In (D), (E), and (F), the metals, metal binding residues and metal-bound water molecules are presented, as well as relevant bond distances.

Closer inspection of the different structures’ metal-binding sites confirms the identity of the previously proposed residues involved in metal binding (Barber-Zucker et al., 2019; Zeytuni et al., 2014b). The Cu^2+^-bound structure contains three binding sites: a central binding site (CS) in which one copper ion is bound by H285 from both monomers and an additional two water molecules, and two ‘symmetric’ peripheral binding sites (PSs), each of which binds one copper ion (Figure 1A, B, D, see metal-ligand distances in Table 3). In both PSs, H264 chelates the copper ion. In one of the PSs, three water molecules appear to stabilise the ion at this location, and one of them bridges between E289 in the adjacent monomer and the metal ion. E268 is also close to the copper ion at this site, however it does not appear to ligate directly or through a water molecule, whereas in the second PS, E268 is much closer to the ion and participates directly in its chelation (Table 3). In this second PS (much like in the first), a water molecule also bridges between E289 in the adjacent monomer and the metal ion. Since E289 has previously been shown to impact MamM function (Zeytuni et al., 2014b), and as binding to the PSs appears to facilitate tighter dimeric packing (Barber-Zucker et al., 2019), we propose that this water molecule plays an important role in the stabilisation of the closed state.

**Table 3:** Coordination distances to ions in MamM CTD metal-bound crystal structures.

|  | E215<br>3<sup>rd</sup>-mer | D249 | H285 | H264 | E289 to<br>water | E268 | Water |
| --- | --- | --- | --- | --- | --- | --- | --- |
| Cu<sup>2+</sup>-<br>bound<br>(6GP6) |  | - | 2.41 <sup>3</sup><br>2.29 <sup>3</sup> | 2.48 <sup>1</sup><br>2.11 <sup>2</sup> | 3.52 <sup>a,1</sup><br>3.92 <sup>b,2</sup> | 2.99 <sup>1</sup><br>3.94 <sup>2</sup> | 3.37 <sup>a,1</sup><br>2.27 <sup>b,2</sup><br>2.88 <sup>2</sup><br>3.83 <sup>2</sup><br>3.51 <sup>3</sup><br>2.54 <sup>3</sup> |
| Cd<sup>2+</sup>-<br>bound<br>(6GMT) | 2.17 <sup>1</sup><br>2.21 <sup>2</sup> | 2.44 <sup>1</sup><br>2.60 <sup>1</sup> | 2.31 <sup>2</sup> | - | - | - | - |
| Ni<sup>2+</sup>-<br>bound<br>(6GMV) | - | - | - | 2.32 | - | - | 2.30<br>2.08<br>2.41<br>2.52 |

All distances were measured using UCSF Chimera (Pettersen et al., 2004).

In the Cd^2+^-bound structure, which contains two bound metals per monomer (i.e. 4 metal sites in total), D249 and H285 each bind one cadmium ion in each monomer. Both cadmium ions are also chelated by E215 from a non-biological-assembly monomer (Figure 1A, C, E and Table 3) and possess half occupancy in the electron density map, suggesting that they may be bound alternately. Mutations of both D249 and H285 to alanine, either separately or together, were previously shown to impact the function of MamM CTD both *in vitro* and *in vivo* (Barber-Zucker et al., 2019; Zeytuni et al., 2014b). This Cd^2+^-bound structural model indicates that both residues are able to bind metals, although this binding alone does not appear sufficient to induce the same conformational change as observed with Cu^2+^. Our previous observation of conformational change toward a closed stable state upon Zn^2+^ binding, was only induced upon metal binding to both the PSs. Hence, binding of Cd^2+^ to the CS alone (in agreement with Zn^2+^ binding to the CS alone (Barber-Zucker et al., 2019)) is not expected to cause a distinct conformational change.

In the Ni^2+^-bound structure, which only contains one bound metal per monomer in the peripheral site, this nickel ion is bound to H264 and to water molecules, possibly implying a non-specific interaction of the Ni^2+^ ion with the histidine (Figure 1A, C, F and Table 3). However, specific chelation to this histidine rather than to other metal-binding residues might suggest a higher affinity for nickel, whilst its location in the protein periphery could also suggest it plays an important role in attracting the metal ions. This proposal is supported by previous studies that demonstrated the impact of H264 on MamM function (Zeytuni et al., 2014b), as well as the new Cu^2+^-bound structure.

Interestingly, all the metal-dependent crystallographic binding sites can be well explained by the metals’ propensities to be bound by specific amino acids: Cu^2+^ has the highest affinity for histidine, followed by Ni^2+^ with Cd^2+^ having the lowest affinity, whilst the aspartate and glutamate binding tendencies show the opposite trend (Barber-Zucker et al., 2017). Overall, the MamM CTD metal-bound crystal structures confirm the previously proposed binding sites and reveal diversity amongst the different metals’ chelation. Since each metal is bound by different residues and only the Cu^2+^-bound structure reveals the apparent conformational change, these crystallographic results would indicate that each metal exhibits a different binding mode and may regulate overall protein function in a distinct manner.

### Dimer packing of the MamM CTD Cu^2+^-bound structure is less tight compared to other CDF CTD bound structures

Previously, a number of CDF protein CTD structures were solved either in their apo state, their Zn^2+^-bound state (YiiP from *Escherichia coli* and YiiP homolog from *Shewanella oneidensis*, EcYiiP and SoYiiP respectively), or in both apo and Zn^2+^-bound states (MamB and CzrB) (Cherezov et al., 2008; Coudray et al., 2013 Lopez-Redondo et al., 2018; Lu et al., 2009; Lu and Fu, 2007; Uebe et al., 2018) (Figure 2A). According to these solved structures, the only protein to exhibit a distinct closure of the V-shape upon metal binding compared with its apo state is CzrB (pdb codes: 3BYP & 3BYR) (Cherezov et al., 2008). The bound state of CzrB exhibits a larger degree-of-closure than that of the MamM CTD Cu^2+^-bound state, demonstrated by the larger differences in the distances and the dihedral angles between equivalent residues in the apo and closed states (Figure 2B, Table 1 and Table 2). This is due not only to a tighter closed state, but also to a slightly more open apo form, compared with those of MamM CTD. MamB, another CDF protein from MTB, exhibits very little conformational change between the apo and Zn^2+^-bound states (Uebe et al., 2018). However, the degree-of-closure in both MamB CTD structures is larger than that of MamM CTD apo structures, and is similar to that of the MamM Cu^2+^-bound structure (pdb codes: 5HO1 & 5HO5; Figure 2C, Table 1 and Table 2). The structures of the entire CDF proteins EcYiiP and SoYiiP (including both the CTD and TMD), have also been solved. In both cases, the CTD is in the bound state (pdb codes: 3H90 & 5VRF, where the CTD structures are highly conserved) (Lopez-Redondo et al., 2018; Lu et al., 2009) and the degree-of-closure is also greater than that of the MamM Cu^2+^-bound state (Figure 2A, Table 1 and Table 2). However, this cannot be directly compared to the overall conformational change exhibited by MamM between apo and bound states, since the apo structures of EcYiip and SoYiiP are not currently available. In conclusion, the MamM CTD Cu^2+^-bound structure shows moderate dimerisation-closure compared with other CDF bound proteins, as was also proposed in our previous study of Zn^2+^-binding and associated conformational change in solution (Barber-Zucker et al., 2019). This may be due to the presence of an additional CTD terminal helix within MamM, which is not found in other CDF protein structures, and which potentially extends the interdimer gap. However, there is variation in the degree-of-closure between the CDF CTD’s of other proteins, suggesting that some degree of flexibility in the CTD conformational changes between the different CDF family members exists. Noticeably, this also depends on the CTD-TMD interface which occurs mainly between the CTD and TMD loops. This interface can vary between the different proteins and thus affect the degree-of-closure exhibited by each individual protein’s CTD.

**Figure 2:**
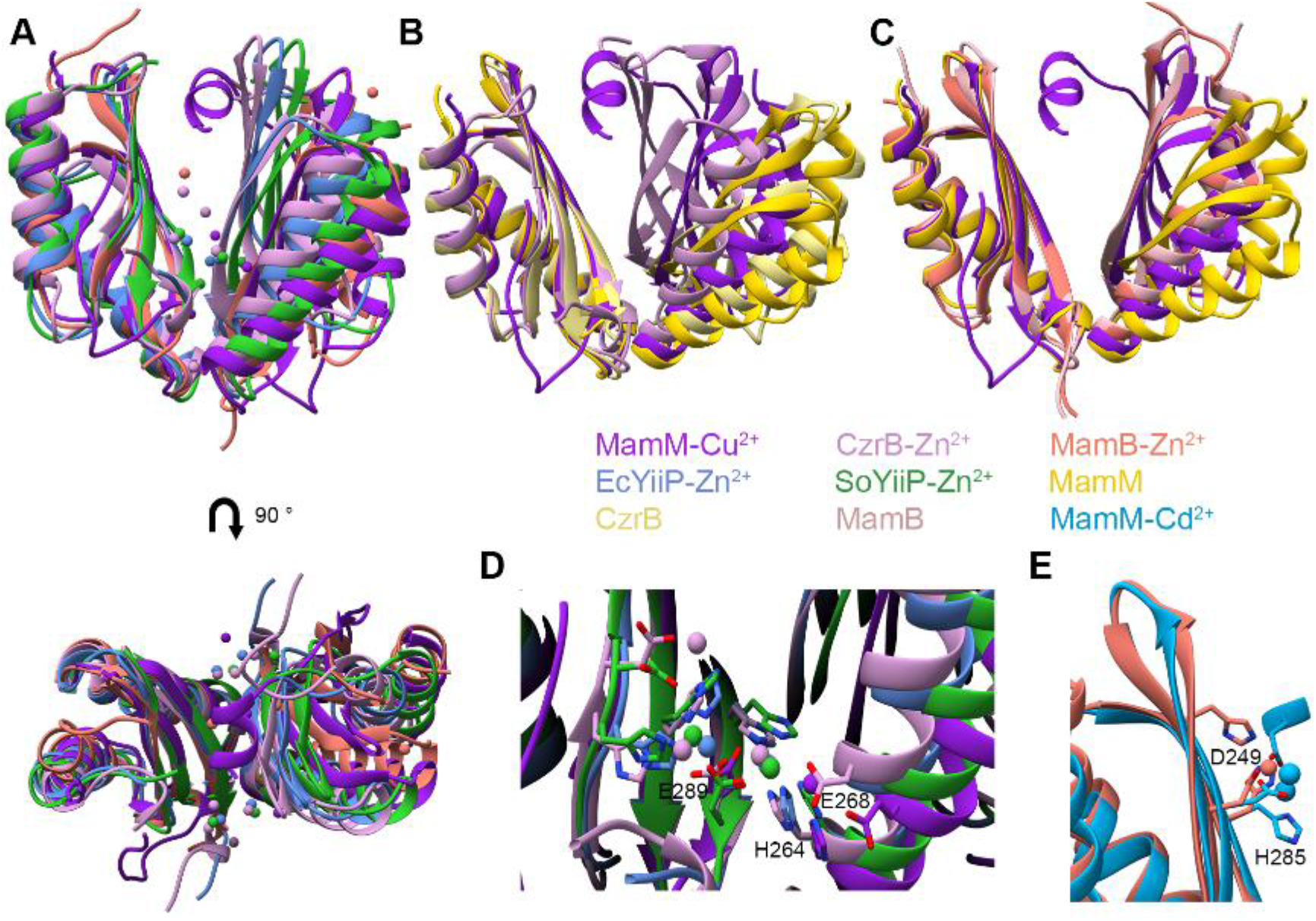
Comparison of the conformational changes and binding sites of different CDFs’ CTDs. Colors legend: MamM Cu^2+^-bound (pdb code: 6GP6, dark purple), CzrB Zn^2+^-bound (Cherezov et al., 2008) (pdb code: 3BYR, light purple), MamB Zn^2+^-bound (Uebe et al., 2018) (pdb code: 5HO1, salmon), EcYiiP Zn^2+^-bound (Lu et al., 2009) (pdb code: 3H90, cornflower blue), SoYiiP Zn^2+^-bound (Lopez-Redondo et al., 2018) (pdb code: 5VRF, green), MamM apo (Zeytuni et al., 2014b) (pdb code: 3W5X, dark yellow), CzrB apo (Cherezov et al., 2008) (pdb code: 3BYP, light yellow), MamB apo (Uebe et al., 2018) (pdb code: 5HO5, light pink), MamM Cd^2+^-bound (pdb code: 6GMT, sky blue). (A) Comparison between all bound CDF CTDs structures at two angles: MamM Cu^2+^-bound, MamB, CzrB, EcYiiP, SoYiiP (all Zn^2+^-bound). Metal cations are presented for each bound structure in the same colors. (B) CzrB exhibits much larger conformational change between apo and bound states, compared to MamM; MamM apo, MamM Cu^2+^-bound, CzrB apo and CzrB Zn^2+^-bound structures are overlapped. (C) MamB exhibits no conformational changes between the bound and apo states, and shows tighter packing compared to MamM apo form; MamM apo, MamM Cu^2+^-bound, MamB apo and MamB Zn^2+^-bound structures are overlapped. (D) Peripheral site binding residues are conserved between MamM, CzrB and YiiP proteins. Metal cations are presented for each bound structure in the same colors; metal-binding residues are presented as sticks. The presented residue labels refer to MamM. (E) Central site binding residues are conserved between MamM and MamB proteins. Metal cations are presented for each bound structure in the same colors; metal-binding residues are presented as sticks. The presented residue labels refer to MamM.

The binding sites present in all MamM CTD metal-bound structures are consistent with those of other CDF protein binding sites. MamM PSs are composed of H264 (Cu^2+^-bound and Ni^2+^-bound structures) and E289 via a bridging water molecule (Cu^2+^-bound structure). In CzrB, six zinc ions are bound to the CTD through three sets of symmetric binding sites (Figure 2D) (Cherezov et al., 2008). Two Zn^2+^ ions are bound in symmetric PSs each by H258 (from one monomer), E282 and H280 (from the second monomer), where residues H258 and E282 are homologous to H264 and E289 in MamM PSs (H280 is homologous to S287 in MamM). Two additional Zn^2+^ ions are symmetrically bound by E282 and additional residues of the same monomer which are not homologous to any of the MamM CTD binding residues, but to residues that do not usually participate in metal binding. These residues also participate in the binding of the last two symmetric Zn^2+^ ions. Overall, this protein-Zn^2+^ interaction network stabilises Zn^2+^ binding in the peripheral sites, which are the only sites to exhibit binding by residues in both monomers, indicating that the PSs are important for conformational change in CrzB. In the YiiP bound structures, two symmetrical pairs of Zn^2+^ ions are bound in the same locations and by homologous residues to the first two Zn^2+^ pairs in CzrB (Figure 2D) (Lopez-Redondo et al., 2018; Lu et al., 2009), further highlighting the importance of the peripheral sites. Although the E289 residue in MamM does not chelate the Cu^2+^ directly in the crystal structure, Zn^2+^ is bound by the homologous residues to E289 in the crystal structures of the other Zn^2+^ bound CDF proteins. Furthermore, E289 is crucial for proper function *in vitro* and *in vivo* (Zeytuni et al., 2014b). Thus, it is clear that E289 in MamM plays an important role in metal-binding for biological function.

The MamM Cu^2+^-bound structure also involves one Cu^2+^ ion bound centrally by H285 from each monomer and an additional two water molecules. Homologous to H285 of MamM is residue H283 in MamB, involved in the chelation of Zn^2+^. In MamB, one Zn^2+^ ion is coordinated in each monomer by residues H283, D247 (homologous to D249 in MamM) and H245 (found in the same location as W247 in MamM) (Figure 2E)^24^. While the MamM Cu^2+^-bound structure involves only H285 – but from both monomers, together binding a single copper ion – the Cd^2+^-bound structure exhibits binding more similar to that of MamB. In the Cd^2+^-bound structure, no cadmium ions are bound to the PSs, and residues of MamB which are homologous to the MamM PSs would not afford effective metal cation binding. Additionally, both metal-bound structures exhibit no conformational changes compared to their apo forms. However, unlike the MamB Zn^2+^-bound structure, Cd^2+^-bound MamM includes two Cd^2+^ ions per monomer (one bound by H285 and the other by D249). Nonetheless, since these Cd^2+^ ions possess only half occupancy in the electron density map, on average just one ion is bound in each monomer, as is the case with MamB. When considering all CDF metal-bound structures, a central site exists only in MamM and MamB, and mutations of these CS residues in both proteins were shown to lead to lower functionality both *in vivo* and *in vitro* (Uebe et al., 2018; Zeytuni et al., 2014b). Noticeably, copper binding in a unique coordination in the CS facilitates subsequent binding to the PS (as was shown for Zn^2+^ (Barber-Zucker et al., 2019)), or vice versa. Overall, the analysis of all known structures suggests that the CS and PSs bind different metals distinctively in MamM, strengthening the previous observation that only when metals are bound to the PSs can the conformational changes effectively occur.

### MamM CTD shows metal-type-dependent change in tryptophan fluorescence signal

Crystal structures often only provide limited and sometimes biased information on the conformation of proteins. Hence, to further characterise MamM CTD binding to different metal cations, we have investigated these interactions in solution. Each MamM CTD monomer contains one tryptophan residue, W247, juxtaposed to the CS. When metal cations bind to MamM CTD, it is proposed that the monomers approach each other, and hence the tryptophan environment will be changed which should lead to a shift in the emission spectrum of the tryptophan residue when excited at the tryptophan-specific wavelength (λ=297 nm). In the case of paramagnetic metals, their binding in close proximity to this tryptophan will cause the tryptophan fluorescence intensity to be quenched. We have shown previously that the addition of Zn^2+^ causes a blue shift in the emission spectrum of MamM CTD, and that the addition of the paramagnetic Fe^2+^ causes signal quenching (Barber-Zucker et al., 2019; Zeytuni et al., 2014b), indicating that MamM CTD binds both cations. In this study we have titrated further metal cations (Cd^2+^, Ni^2+^, and Cu^2+^, in which their bound crystal structures were detected, as well as Mn^2+^, which was shown to be transported by numerous CDF proteins (Barber-Zucker et al., 2017; Cubillas et al., 2013)) into MamM CTD solution samples and measured the concentration-dependent changes in the tryptophan emission spectra (ratios of 0:1 to 100:1 metal:protein, Figure 3). As shown previously for Zn^2+^ (Figure 3A, adopted from (Barber-Zucker et al., 2019)), in the case of the diamagnetic Cd^2+^, a shift in maximum wavelength could be observed (Figure 3B), suggesting that the binding of Cd^2+^ leads to a change in the tryptophan environment. However, whilst the spectral shift is accompanied by an increase in intensity for Zn^2+^, the intensity decreases when Cd^2+^ is added and the blue shift is smaller (Figure 3F, G). The Cd^2+^-bound crystal structure suggests no conformational change compared to the apo form, whilst it was previously shown that the addition of Zn^2+^ causes a tight closure of the CTD (Barber-Zucker et al., 2019). Since cadmium is much bigger in size than zinc, Cd^2+^ binding might restrict the ability of the dimer to pack tighter, which could result in a smaller shift and the difference in the fluorescence intensity observed for Cd^2+^.

**Figure 3:**
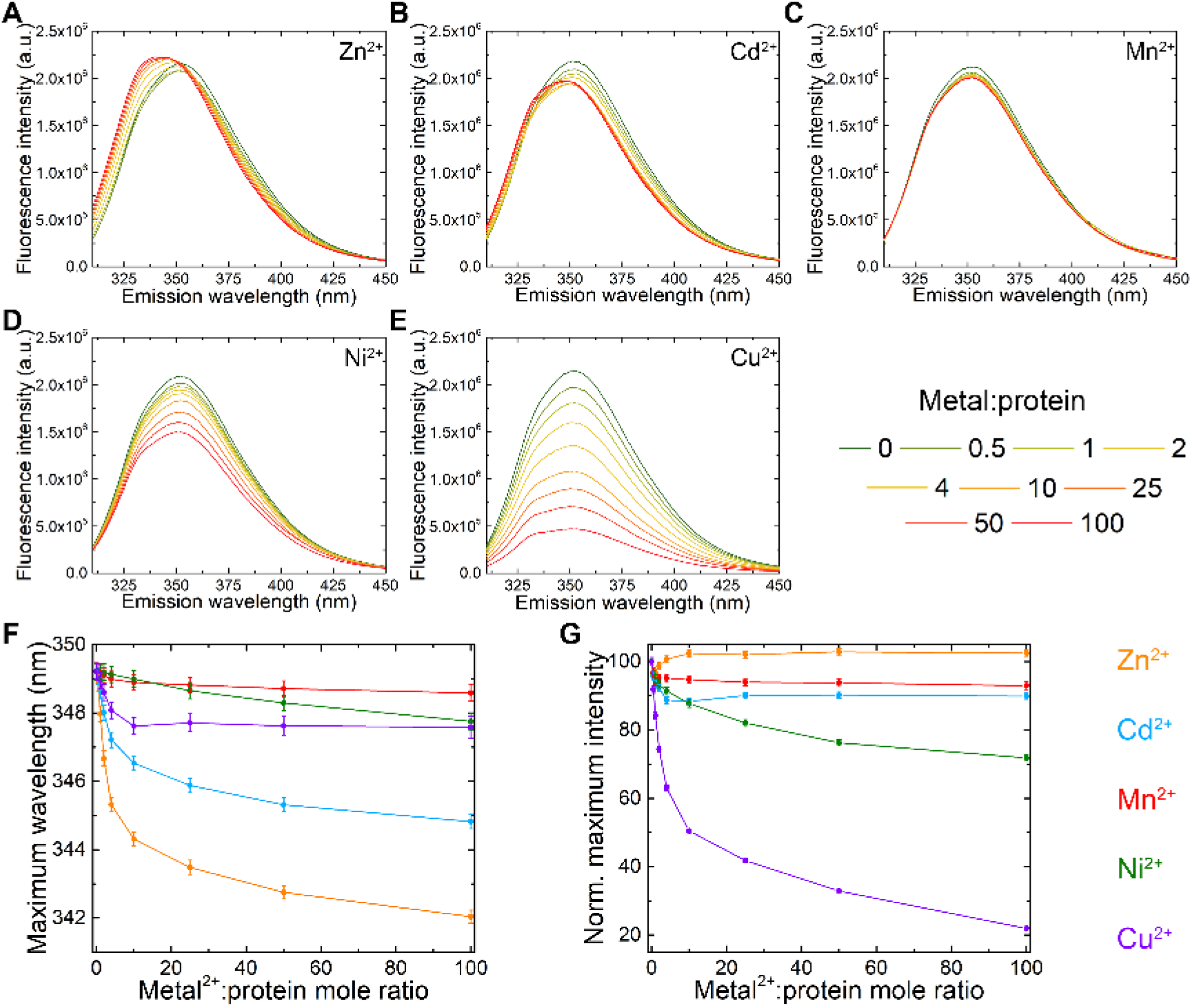
Fluorescence spectral scans of MamM CTD with different metals. All emission spectra of 5 μM MamM CTD were titrated using different metal solutions to reach different MamM CTD:Metal^2+^ ratios (intensities were normalised due to the change in MamM CTD concentration). Samples were measured at λ_ex_ 297 nm, and the emission spectrum for each metal concentration was recorded at 310-450 nm. Each presented spectrum is the average of three independent measurements. (A) Emission spectra of WT with different Zn^2+^ concentrations (adopted from (Barber-Zucker et al., 2019)), (B) Emission spectra of WT with different Cd^2+^ concentrations, (C) Emission spectra of MamM CTD with different Mn^2+^ concentrations, (D) Emission spectra of MamM CTD with different Ni^2+^ concentrations, (E) Emission spectra of MamM CTD with different Cu^2+^ concentrations, (F) λ_max_ as function of Metal^2+^: MamM CTD mole ratio, (G)Normalised fluorescence intensity compared to no Metal^2+^ as function of Metal^2+^: MamM CTD mole ratio. In (F) and (G), Zn^2+^ is presented in orange, Cd^2+^ in blue, Mn^2+^ in red, Ni^2+^ in green and Cu^2+^ in purple, and error bars are shown within the symbols.

Alternatively, it is also possible that the Cd^2+^ ions, which are bound in close proximity to W247, simply influence the tryptophan fluorescence signal differently. Mn^2+^, Ni^2+^ and Cu^2+^ are all paramagnetic, hence their binding to the CTD would be expected to cause observable fluorescence quenching leading to a stronger signal than their binding-dependent spectral shift. However, the addition of Mn^2+^ only reveals minor changes in the fluorescence intensity (Figure 3C, G) and in the maximum wavelength (Figure 3F), suggesting that Mn^2+^ cannot bind to the CTD in a way that would cause a change in the fluorescence spectra. In contrast, Ni^2+^ and Cu^2+^ demonstrate clear metal-concentration-dependent quenching (Figure 3D, E, G) and a slight spectral shift (Figure 3F), with Cu^2+^ exhibiting stronger quenching and a larger shift. Overall, these results indicate that MamM CTD can bind all the examined metals with the exception of Mn^2+^.

### MamM CTD exhibits metal-type-dependent binding thermodynamics

In order to accurately calculate the affinity of each metal to MamM CTD, other thermodynamic parameters, and the number of metal cations that can bind to the CTD, ITC measurements were performed (Figure 4). Here Cd^2+^, Ni^2+^, Cu^2+^ and Mn^2+^ were titrated into MamM CTD solution samples. All ITC parameters are summarised in Table 4, including those for previously titrated Zn^2+^ (Barber-Zucker et al., 2019) (note: metal binding to MamM CTD requires a slightly basic pH due to the metal chelation by histidine residues, therefore titration of Fe^2+^ leads to iron precipitation and hence its binding parameters cannot be accurately determined). ITC results indicate that all metals except Mn^2+^ induce a heat change when titrated into a MamM CTD solution. Hence taken together with the fact that Mn^2+^ revealed no fluorescence shift, we propose that MamM CTD is unable to bind Mn^2+^.

**Table 4:** Thermodynamic parameters of MamM CTD binding to different metals.

|  | Zn<sup>2+</sup> <sup>a</sup> | Cd<sup>2+</sup> | Ni<sup>2+</sup> | Cu<sup>2+</sup> |
| --- | --- | --- | --- | --- |
| K<sub>d</sub> (μM) | 31.50±3.40 | 48.39±2.61 | 17.71±3.35 | 6.00±3.83 |
| N | 2.80±0.24 | 1.23±0.08 | 3.26±0.19 | 1.56±0.15 |
| ΔH (kJ mol<sup>-1</sup>) | -4.20±0.10 | -13.08±1.56 | -7.78±2.38 | -71.80±23.59 |
| ΔG (kJ mol<sup>-1</sup>) | -25.71±0.26 | -24.64±0.13 | -27.17±0.48 | -30.13±1.59 |
| ΔS (J mol<sup>-1</sup> K<sup>-1</sup>) | 72.15±0.98 | 38.76±4.81 | 65.03±9.59 | -136.72±83.76 |

ITC data were fitted to independent + blank (constant) models in the TA NanoAnalyze Data Analysis software.
Each parameter represents three independent measurements after control reduction (metal at the same concentration titrated into buffer solution).
The buffer used for all titrations: 10 mM Tris-HCl pH 8.0 and 150 mM NaCl.
<sup>a</sup> The data are taken from (Barber-Zucker et al., 2019).

**Figure 4:**
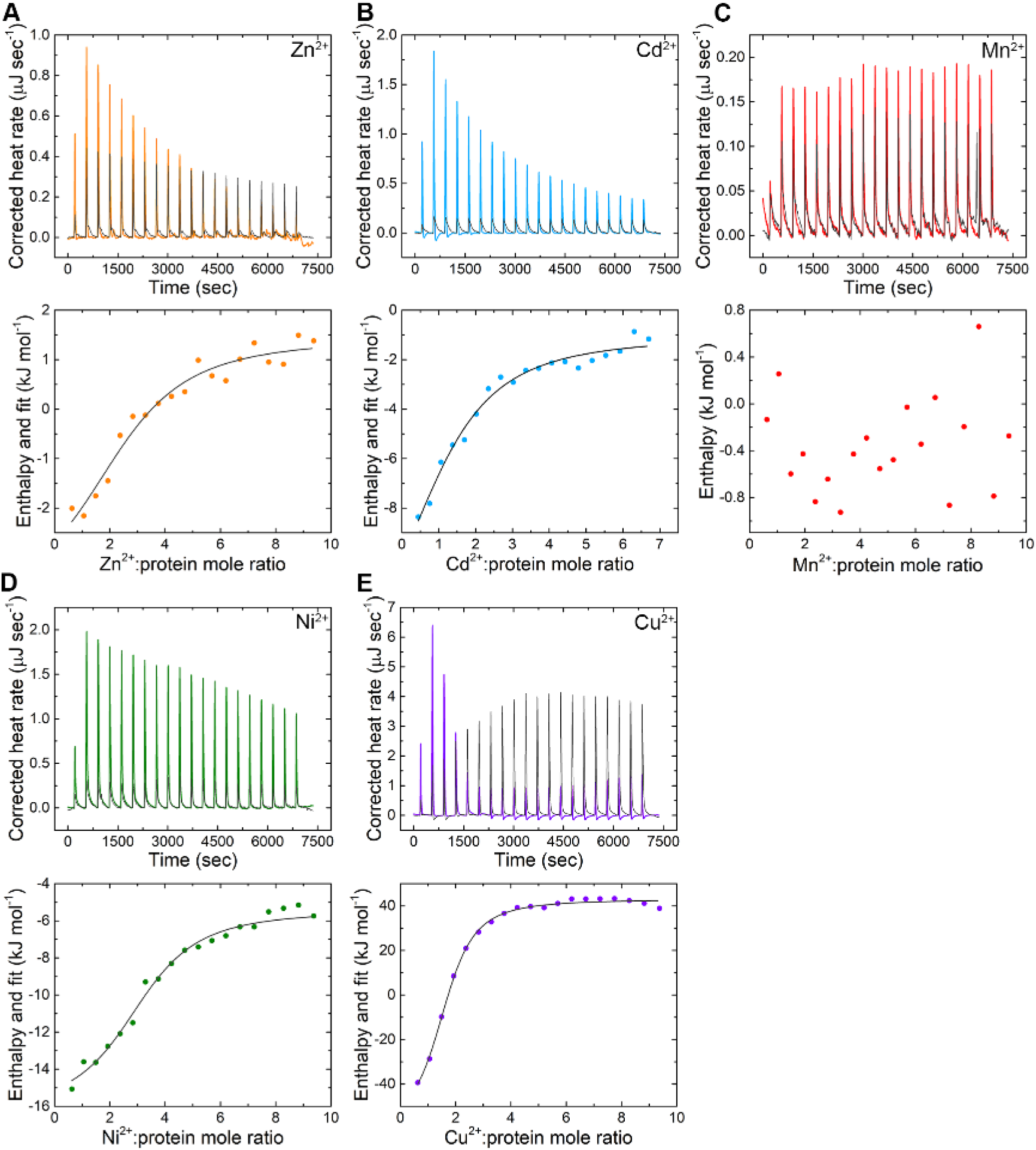
ITC titration measurements of MamM CTD-metals interactions. (A) with Zn^2+^ (adopted from (Barber-Zucker et al., 2019)), (B) with Cd^2+^, (C) with Mn^2+^, (D) with Ni^2+^, (E) with Cu^2+^. Top panels: Corrected heat rate curves for a representative titration of MamM CTD with each metal as a function of time. For each metal, the heat flow curve of the control (metal titration into buffer, dark gray) is given as a reference. Bottom panels: Titration data after peak integration as a function of Metal^2+^: MamM CTD mole ratio, and data fitting for all metals except Mn^2+^. See Table 4 for binding parameters.

The K_d_ values determined vary from 6-48 μM, with Cu^2+^ having the greatest affinity followed by Ni^2+^, Zn^2+^ and Cd^2+^ (Table 4). As revealed in all metal-bound crystal structures and previous *in vivo* studies (Zeytuni et al., 2014b), histidine residues are the major components for metal binding. Again the larger size of the Cd^2+^ ion may explain its low affinity as its coordination may be less preferred, however a lower propensity to bind histidine residues compared to the other metals (Barber-Zucker et al., 2017) may also result in a lower affinity to proteins with multiple histidine-containing binding sites such as MamM CTD. Cu^2+^ has the highest propensity to bind histidine, which clearly explains its high affinity to MamM. Ni^2+^ also exhibits a high propensity for histidine binding compared to Zn^2+^ and Mn^2+^ (Barber-Zucker et al., 2017). Although all binding reactions are exothermic (Table 4), Cu^2+^-binding displays much lower enthalpy, which means it is favoured in terms of the heat release. Moreover, while Ni^2+^, Zn^2+^ but also Cd^2+^ display high entropy, the entropy of Cu^2+^ binding is negative, suggesting that its binding results in a more ordered state of the cations compared to its free state in solution. Summarizing all of the above, the negative Gibbs free energy of the binding reactions imply that all metal-binding reactions are preferred to the apo state (excluding Mn^2+^), yet MamM CTD shows clear preferences for the binding of specific metals over others.

The numbers of metal cations bound per dimer is also given by the ITC measurements, which varies significantly between metals (Table 4). For Zn^2+^ and Ni^2+^, ~ 3 cations are bound per dimer, in agreement with the Cu^2+^-bound structure. The Ni^2+^-bound crystal structure exhibits 2 cations per dimer in the PSs, however it shows no conformational changes when compared with the apo state and binding appears to be non-specific - disagreements between the metal-bound structure and ITC results can therefore be expected.

For Cd^2+^ and Cu^2+^ lower number of metal cations bound per dimer is observed: 1.23 ± 0.08 for Cd^2+^ and 1.56 ± 0.15 for Cu^2+^. The number of metals observed in metal-bound structures are clearly not in agreement with these ITC observations, which are much lower. The Cd^2+^-bound structure reveals four metal binding sites per dimer; however, the low occupancy observed in the electron density map would suggest that actually only two cations are bound at any one time. Moreover, binding in the crystal structure is mediated by a glutamate residue from a third monomer (from an adjacent unit cell) which may simply be a crystallographic artefact not necessarily expected to be observed in solution. Nevertheless, although the tryptophan-fluorescence scans suggest otherwise, this metal-bound structure does not exhibit any conformational change as also observed for the Ni^2+^-bound structure. Hence, the ~ 1 Cd^2+^ ion bound per dimer determined from the ITC results suggests that binding at the CS is of a different nature to that displayed in the crystal structure.

The unexpectedly low number of Cu^2+^ ions bound in solution, as compared to the crystal structure, could have several explanations: it is quite possible that Cu^2+^ binding to the CS is silent in terms of the heat change (as was also shown previously for Zn^2+^-binding to the CS (Barber-Zucker et al., 2019)), hence the thermodynamic parameters may relate only to the binding of two ions to the PSs (1.56 ~ 2, and thus three Cu ions in total); alternatively, in solution only one ion might bind at the CS, and none at the PSs (then 1.56 ~ 1, in clear disagreement with the structure); however the opposite may also be true, in which two ions are bound to the PSs, and none to the CS (then again 1.56 ~ 2, and thus two overall, again in disagreement with the structural data). Since binding to the PSs has previously been shown to be the only cause of conformational change in MamM and other CDF CTDs, one would expect that if Cu^2+^ only binds to the CS, then no conformational changes should occur in solution, and that if Cu^2+^ only binds to the PSs, then a tighter conformation compared to the apo form would be detected in solution. The large tryptophan-fluorescence signal quenching demonstrated by titration of Cu^2+^ into MamM CTD indicates that the copper ions are bound in close proximity to W247, hinting at an occupied CS. However, the tryptophan-fluorescence resolution is not sufficient to distinguish between these proposed options.

### EPR and PELDOR spectroscopy observes metal-bound MamM CTD conformational changes

To probe divalent metal binding and the associated conformational changes reflected by intra-molecular distance, site-directed spin labelling (SDSL) was employed in combination with both continuous-wave electron paramagnetic resonance (cw-EPR) and pulsed electron-electron double resonance (PELDOR) spectroscopic studies (Barber-Zucker et al., 2019). These methods provide a wealth of dynamic and structural information, via observation both of changes in the local environment of an attached exogenous spin label elucidated by cw-EPR, and the determination of long-range (typically between 2-8 nm) distance measurements and distance distributions afforded by PELDOR spectroscopy (Mullen et al., 2016). For SDSL, a stable paramagnetic (1-oxyl-2,2,5,5-tetramethylpyrrolidin-3-yl)methylthiosulfonate spin label (MTSL) reacts specifically with cysteine residues present within the protein, hence allowing EPR to report on the protein via these specifically labelled sites. We have previously reported (Barber-Zucker et al., 2019) that using only one of the two intrinsic MamM CTD cysteine residues simplifies EPR measurements, hence the studies reported here use a C267S variant of MamM CTD, in which only the C275 site available for labelling. *In silico* predictions using Multiscale Modelling of Macromolecules (MMM), a Matlab^©^-based open-source modelling toolbox (Jeschke, 2018), and the apo WT crystal structure (PDB code: 3W5Y) predict an average inter-spin distance of 4.0 nm between the labelled cysteines (C275) in each protomer (Barber-Zucker et al., 2019). This result is replicated with the C267S crystal structure (PDB code: 6G55) (Figure 5A).

**Figure 5:**
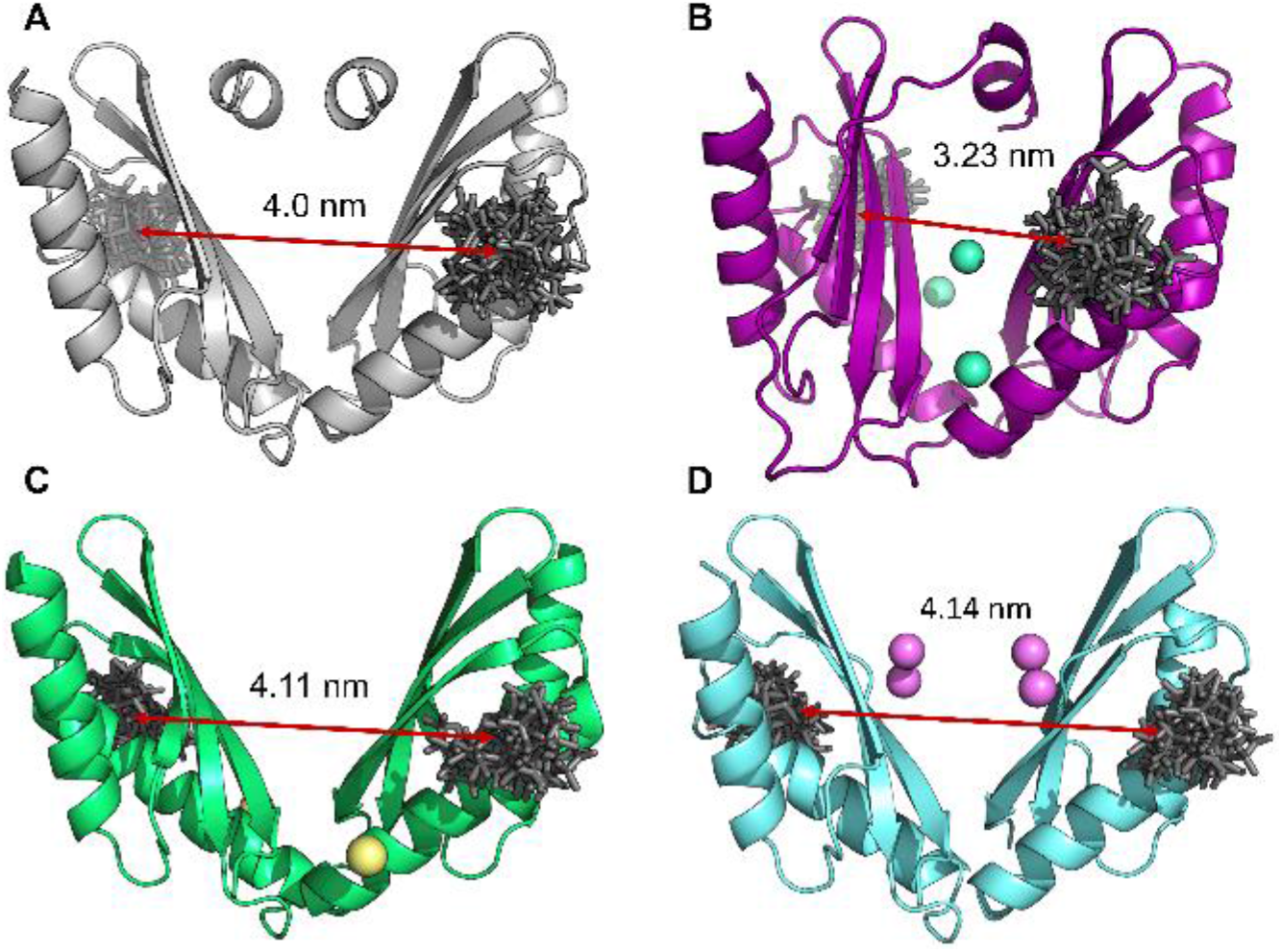
MamM CTD structural models built using the published crystal structures. Cartoon representation of (A) dimeric apo MamM CTD C267S (PDB code: 6G55) (Barber-Zucker et al., 2019), (B) Cu^2+^-bound MamM CTD (PDB code: 6GP6), (C) Ni^2+^-bound MamM CTD (PDB code: 6GMV) and Cd^2+^-bound MamM CTD (PDB code: 6GMT) with the calculated rotamer libraries of MTSL attached to position C275 shown as sticks, and with metal ions shown as spheres. *In silico* modelling of MTSL rotamer libraries and calculation of the average inter-spin distance was performed using Multiscale Modelling of Macromolecules (MMM)(Jeschke, 2018), a Matlab^©^-based open-source modelling toolbox.

cw-EPR studies at X-band of complex dynamic proteins affords the observation of changes in the local environment of an attached exogeneous spin label, as well as the overall mobility of the protein in different states. Studies have been undertaken to ascertain what effect, if any, the addition of zinc and copper metals (to which the protein has a lower and higher affinity respectively) has on the local spin label environment of C267S_SL_. Indeed, cw-EPR spectra of these metal-bound C267S_SL_ sample differs substantially from that of the apo protein (Figure 6A). This is indicative of an altered local environment around the spin label with both metals when compared to the apo state. The dynamics exhibited by spin labels attached to a protein in the solution state are described by the rotational correlation time, which is made up of three factors: (1) the rotational correlation time of the protein itself; (2) the backbone motions of the protein at the point of spin label attachment; and (3) the intrinsic motion of the spin label itself. Line shapes of continuous wave spectra at X-band, and in particular their broadness, are dominated by the latter two of these contributions. Binding of metal ions to the protein resulting in conformational change will affect these modes of motion, hence resulting in different spectral line shapes observed.

**Figure 6:**
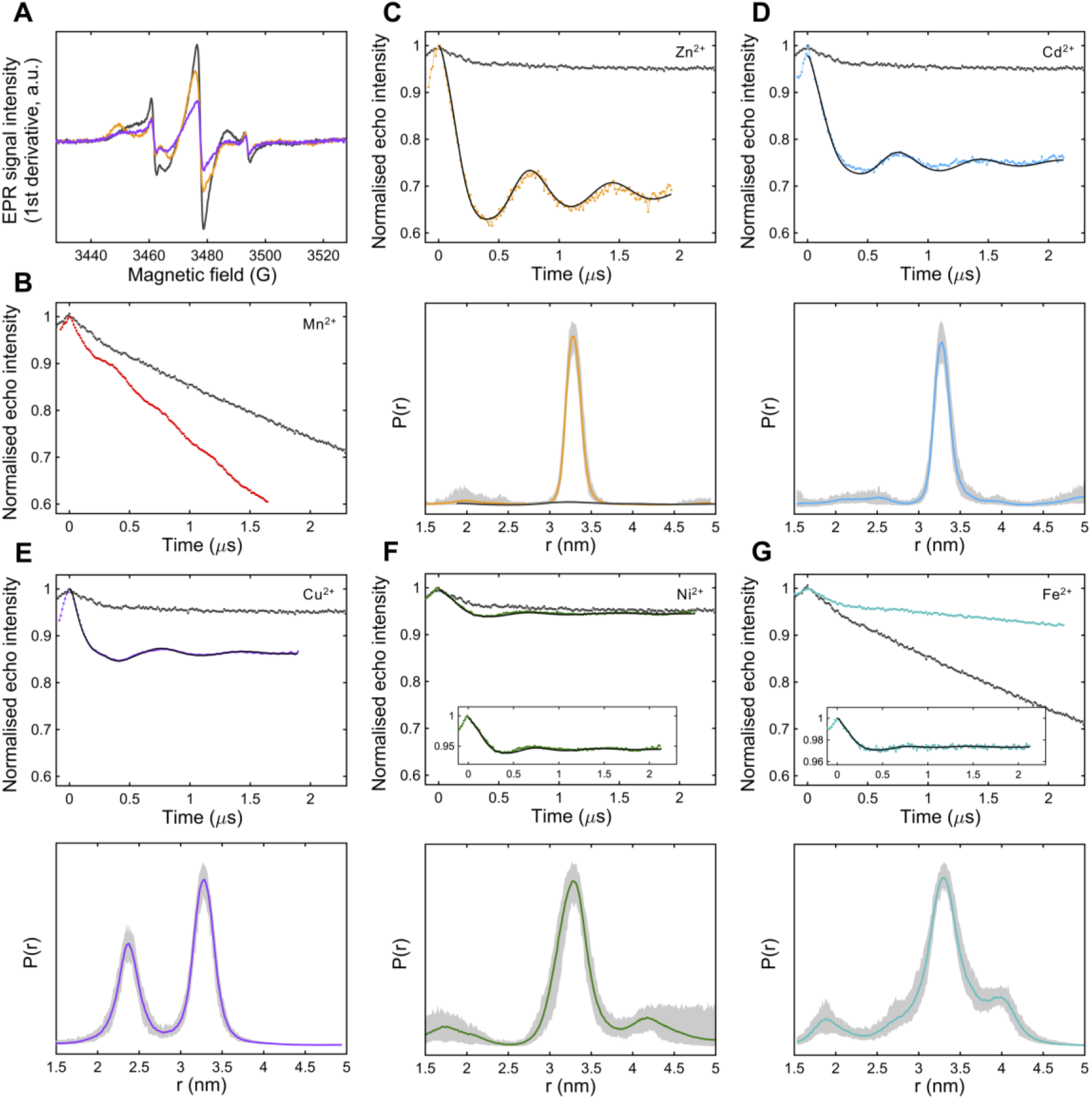
Room temperature continuous-wave EPR measurements (A) performed on apo MamM C267S_SL_ (grey), C267S_SL_-Zn^2+^ (orange) and C267S_SL_-Cu^2+^ (purple) revealing quenching of the EPR signal intensity upon addition of Cu^2+^ as compared to both the apo and Zn^2+^-bound states. (B-G) Four-pulse PELDOR experiments of doubly spin-labelled MamM CTD C267S, containing C275 as the labelling position, for the measurement of inter-spin label distances. Results were obtained for samples incubated with a range of divalent transition metal ions, including Mn2+ (B), Zn^2+^ (C, adopted from (Barber-Zucker et al., 2019)), Cd^2+^ (D), Cu^2+^ (E), Ni^2+^ (F), and Fe^2+^ (G). Top panel: The normalised background-corrected dipolar evolution of metal-bound samples (bright colours) presented together with the apo spin labelled protein (grey). Exceptions; For both Mn^2+^ and Fe^2+^ the non-baseline corrected time traces are shown to clearly demonstrate the change in decay caused by specific versus non-specific metal binding. Bottom panel: Normalised distance distributions (where appropriate) of metal bound C267S_SL_. The distance distribution of apo C267S_SL_ is also shown for reference, plotted against the Zn^2+^-bound results and shown in dark grey. Standard deviations in the form of 95 % confidence intervals are also given in each case and represented by light grey areas around the lines. Fitting of all dipolar evolutions and subsequent distance distributions were calculated with DEERNet (Worswick et al., 2018).

Our previous PELDOR studies of spin labelled C267S MamM CTD (C267S_SL_), undertaken on apo and Zn^2+^-bound forms of the protein, afforded a direct comparison of the inter-spin distances giving insight into the conformational changes associated with Zn^2+^ binding (Barber-Zucker et al., 2019). Hence the 4.0 nm average distance reported from *in silico* predictions above for the apo form is rather misleading since these PELDOR results clearly indicate that in the apo state no single defined conformation is adopted. Only upon addition of Zn^2+^ (forming C267S_SL_-Zn^2+^) is there a clear change in the overall conformation of the protein (Figure 6C). A very narrow distance distribution for C267S_SL_-Zn^2+^ with a mean distance of 3.3 nm was interpreted as a change from a broad dynamic state of apo C267S, to a single closed conformation upon Zn^2+^-binding. Further, stoichiometric Zn^2+^ titrations were followed by PELDOR and suggested a maximum binding of 3-4 Zn^2+^ ions per dimer, in good agreement with ITC experiments (Barber-Zucker et al., 2019).

Here we have undertaken a further cw-EPR and PELDOR study on C267S_SL_ incubated with a range of divalent metal cations, including Cd^2+^, Cu^2+^, Ni^2+^, Mn^2+^ and Fe^2+^ (the natural substrate of MamM). Overall our results indicate that whilst most PELDOR traces and associated distance distributions in the presence of divalent metal ions do show distinct differences from the apo protein, not all metal ion effects are equivalent.

The addition of CdCl_2_ to C267S_SL_ (x5 excess, forming C267S_SL_-Cd^2+^), results in a distinct change in the PELDOR time trace obtained as compared with the apo case. The single and quite narrow distance distribution is predominant with an average distance of 3.3 nm (Figure 6D) and is very similar to the distribution obtained previously with Zn^2+^ (Figure 6C, adopted from (Barber-Zucker et al., 2019)). This would also suggest a closure of the V-shaped dimer upon Cd^2+^ binding, resulting in a single metal-bound conformation from a largely dynamic apo state. While the nature of this binding appears to be similar to that of Zn^2+^-bound MamM CTD, the Cd^2+^-bound crystal structure model does not suggest such metal-dependent conformational change. Obviously the MMM inter spin distance prediction, which uses the crystal structure PDB coordinates (Figure 5D), is 4.14 nm, and is thus similar to that predicted for the apo protein. Since the occupancy for each metal binding site in the solved crystal structure is rather low (0.5), and taken together with ITC data that approximately only one Cd^2+^ ion is bound per dimer as well as the observed change in tryptophan fluorescence, this would suggest that in solution, only one Cd^2+^ binds at the CS. This is in contrast to Zn^2+^ binding since it appears that only metal binding at the CS is required to induce the conformational change. This is possible due the larger size of the Cd^2+^ ion which, once bound to the CS residues of just one monomer, is in close enough proximity to attract the CS binding residues of the second monomer, forcing the observed conformational change.

The addition of paramagnetic Mn^2+^ (x5 excess) to C267S_SL_ does not result in a change in the dipolar evolution time trace as seen when other divalent metals are added (Figure 6B). There are no periodic oscillations arising from dipolar coupling between the two nitroxide moieties, but simply only a slightly modified decay function modulated by the presence of deuterate glycerol (which is not present when protonated glycerol is used, data not shown). Thus, PELDOR taken together with the tryptophan fluorescence and ITC data provide clear evidence of the inability of MamM CTD to bind Mn^2+^. Mn^2+^ has a greater affinity to magnetite, both when abiotically synthesised or in magnetotactic bacteria as compared to other transition metal cations (Amor et al., 2015). Furthermore, uncultivated magnetotactic bacteria, which are exposed to high concentrations of manganese, can incorporate it into the iron-based magnetic particles (Keim et al., 2009). Based on our results and these observations we propose that in order to avoid overflow of manganese in the magnetosomes and a sequential incorporation of it into magnetite, MTB have developed a CTD-related mechanism for discrimination against manganese.

Incubation of C267S_SL_ with NiCl_2_ (x5 excess) forming C267S_SL_-Ni^2+^, however, results in a distinct change in the PELDOR trace as compared to the apo protein (Figure 6F). Again, the resulting distance distribution reveals a single peak centred at 3.3 nm analogous to both Zn^2+^-and Cd^2+^-bound MamM CTDs and is indicative of a single V-shaped conformation. This, however, is in disagreement with the solved Ni^2+^-bound crystal structure of MamM CTD, which revealed no conformational change in either its monomeric fold or dimeric structure when compared to the apo form (MMM results in Figure 5C). If Ni^2+^-binding in the crystal structure is, however, non-specific, as suggested by the metal-ligand variations within the solved structure, then metal-dependent conformational changes in solution in the presence of Ni^2+^ are still be possible. Nonetheless, the shallower modulation depth (Figure 6F) might suggest that a smaller fraction of nitroxide spins are actually dipolar coupled and thus interacting. ITC results indicate binding of 3 Ni^2+^ ions per dimeric protein with mild affinity compared to the other metals, and the observation of tryptophan fluorescence quenching in the presence of Ni^2+^ would imply Ni^2+^ binding either at or close to the central site. Taken together, and considering that the nickel ion’s radius is comparable to that of a zinc ion, we propose that Ni^2+^ binds both to the CS (one ion) and the PSs (two ions), with binding to the latter being associated with a conformational change similar to that observed for Zn^2^ ^+^.

The room temperature cw-EPR experiments of C267S_SL_ incubated with an excess of Cu^2+^ (forming C267S_SL_-Cu^2+^) suggest a partial quenching of the EPR signal as compared to both the Zn^2+^ bound and apo states (Figure 6A). PELDOR studies of C267S_SL_-Cu^2+^ to ascertain whether or not conformational change also occurs in the presence of Cu^2+^ were undertaken using various ratios of Cu^2+^ to protein. However, Cu^2+^ ions are EPR active (S = 1/2) giving EPR signals which overlap with that of the spin label, thus this excess was chosen to be equal to the number of proposed binding sites to avoid any exogenous, spurious Cu^2+^ EPR signals arising from non-specifically bound copper. PELDOR is a two-frequency experiment in which microwave frequencies are selected which allow the pumping and detection of specific spins. PELDOR of C267S_SL_-Cu^2+^ can be conducted in the same manner as previously (i.e. between the two spin labels) but may also be used to probe Cu^2+^ -Cu^2+^ and Cu^2+^ -SL distances as well. The MMM analysis of the Cu^2+^-bound structure (Figure 5B) predicts an inter-spin label distance of 3.23 nm while the resultant distance distribution of the experimental PELDOR trace reveals a bimodal distribution with two mean distances of 2.4 and 3.24 nm (Figure 6E). At first glance this could imply two different inter-spin label (SL-SL) distances, however since the Cu^2+^ species overlaps with the spin label in the EPR spectrum, this PELDOR trace may not only comprise SL-SL distances but in addition, we may have contributions from SL-Cu^2+^ and Cu^2+^ - Cu^2+^ distances. The 3.24 nm distance agrees well with both the predicted MMM SL-SL distance, as well as other metal-bound PELDOR results, and hence we propose that this corresponds to the SL-SL distance again indicative of a single closed V-shaped conformation. The second observed distance at 2.4 nm could correspond to either a Cu^2+^-SL or Cu^2+^-Cu^2+^ distance. Considering the fluorescence data which indicates binding of Cu^2+^ in the CS, and previous PELDOR measurements in which conformational change was shown to be associated with metal binding to the PSs (Barber-Zucker et al., 2019), we propose that binding to the CS is silent in terms of heat change and that the Cu^2+^-bound crystal structure represents well both the conformation in solution and the copper binding sites.

Current experimental evidence for Fe^2+^ binding to MamM CTD, both via biophysical and structural methods, is very limited. This is due largely to the difficulty associated with preparation of bound Fe^2+^ samples; when working at basic pH, Fe^2+^ is readily oxidised to Fe^3+^, which notoriously forms insoluble complexes. Nevertheless, it proved possible to prepare Fe^2+^ incubated sample using argon gas to largely eliminate any oxygen present with C267S_SL_ and an excess of Fe^2+^ (x5 excess, forming C267S_SL_-Fe^2+^). Whilst there was slight indication of Fe^3+^ precipitation, PELDOR yielded results clearly different from the apo state: non baseline-corrected, time-domain data are shown to indicate these differences more clearly and a single distance distribution with a mean distance of 3.3 nm is obtained (Figure 6G). Despite the low modulation depth, probably due to very low levels of Fe^2+^ binding, the difference to the apo state is quite apparent, providing good indication of a more rigid structure although the distance distribution is slightly broader than for Zn^2+^/ Ni^2+^/Cu^2+^/Cd^2+^. Interestingly, the sample became slightly yellow in colour upon incubation with the metal salt, which could not be reproduced upon incubation of the buffer alone with the same concentration of salt. It is therefore likely that the protein also catalyses the iron oxidation process resulting in formation of unbound (or non-specifically bound) yellow Fe^3+^ complexes. More importantly, this is the first high-resolution structural observation of Fe^2+^-binding to MamM CTD, and these results indicate that studies with other metals (including Zn^2+^ for which substantial work has been done previously) do reflect biologically-relevant states of the protein, since all the metal-bound proteins appear to adopt a similar conformation in solution, which is most likely similar to the crystal Cu^2+^ - bound conformation.

### Concluding remarks

The participation of CDF protein CTDs in metal selectivity was previously studied at the cellular level. Here, for the first time, structural characterisation of a CDF protein CTD in the presence of various metals reveals the direct effect of different metals on the CTD, with each metal exhibiting distinct binding parameters, sites and conformation. This study uniquely includes high resolution structural data in both the crystalline form as well as in solution which are complementary to one another, and in combination exhibit the differences between the binding modes to the native metal and other metals – indicative of a metal selectivity mechanism whereby manganese is discriminated against. Furthermore, these results present slight differences in the degree-of-closure between the apo and bound CTDs amongst different CDF proteins, indicating that it depends not only on the protein in study but also on the metal identity.

## Acknowledgements

We thank Dr. Anat Shahar for her help with the crystallographic data collection and analysis. SBZ and RZ are supported by the Israel Ministry of Science, Technology and Space, the Israel Science Foundation (grant no. 167/16) the European Molecular Biology Organization and the EU (CMST COST Action CM1306 Understanding Movement and Mechanism in Molecular Machines). JH was supported by the UKRI Biotechnology and Biological Sciences Research Council Norwich Research Park Biosciences Doctoral Training Partnership (Grant number BB/M011216/1). FM was supported in part by the Royal Society (FM was a Wolfson Research Merit Award Holder). JH and FM are also supported by the the EU (CMST COST Action CM1306 Understanding Movement and Mechanism in Molecular Machines).

## Author Contributions

All authors designed the study, analyzed the results and wrote the paper. SBZ, JH and AF conducted the experiments. All authors approved the final version of the manuscript.

## Methods

### Expression and purification

*mamM* CTD gene (UniProt Q6NE57 residues 215-318) was previously cloned into pET28a(+) vector (Novagen, Merck Biosciences, Germany) (Zeytuni et al., 2012). MamM CTD was expressed and purified as previously described for MamM CTD M250L mutant (Barber-Zucker et al., 2016b; Zeytuni et al., 2012). The C267S mutation for PELDOR studies was applied to the pET28a-MamM-CTD vector, expressed and purified as previously described (Barber-Zucker et al., 2019). For all experiments, protein concentration was determined by measuring protein absorption at 280 nm.

### Crystallisation and structure determination

20 mg mL^−1^ purified MamM CTD in buffer containing 10 mM Tris pH 8.0, 150 mM NaCl, 5 mM β-mercaptoethanol, and either 3.375 mM CuSO_4_ (Cu^2+^-bound structure), 3.375 mM CdCl_2_ (Cd^2+^-bound structure) or 3.375 mM NiCl_2_ (Ni^2+^-bound structure), was crystallised using the vapor diffusion method at 293 K (0.3 μL protein with 0.3 μL reservoir solution for all protein-metal pairs). Crystals were harvested with or without treatment of cryo-agent and flash-frozen in liquid nitrogen. Data collection was performed on a single crystal at a temperature of 100 K. All cryo conditions, data collection details, data reduction and scaling, phasing and refinement details are given in Tables S1-S3.

### Least-squares overlaps

All CDFs’ CTD structures were overlapped and RMSDs were calculated using the iterative magic fit tool or the domain fragment alternate fit tool of Swiss-PdbViewer 4.1.0 (Guex and Peitsch, 1997). MamM CTD structures and overlapped structures’ figures were prepared using UCSF Chimera package, version 1.12 (Pettersen et al., 2004) or PyMOL, version 2.0.6 (PyMOL Molecular Graphics System, Schrödinger, LLC).

### Fluorescence spectrometry

Changes in tryptophan intrinsic fluorescence were recorded using Fluorolog^®^-3 (HORIBA Scientific, Edison, NJ, USA) equipped with quartz cell with 1 cm optical path length at ambient temperature. Samples of 1 mL MamM CTD at 5 μM concentration in buffer A (containing 10 mM Tris pH 8.0, 150 mM NaCl) were titrated using 2.5 mM metal solution (CdCl_2_, NiCl_2_, MnCl_2_, CuSO_4_) in the same buffer to reach different concentrations. Samples were measured at λ_ex_ 297 nm, and the emission spectrum for each metal concentration was recorded at 310– 450 nm. For each metal, the titration was replicated three times, and each spectrum was fitted to Extreme function by OriginPro (R-Square (COD) > 0.98) (OriginLab Corporation, Northampton, MA, USA). The maximum wavelength (wavelength at maximum intensity) and the intensity at that wavelength (maximum intensity) were averaged for each metal concentration. Errors are reported as the standard deviation.

### Isothermal titration calorimetry

ITC measurements were performed in a low-volume Nano ITC calorimeter (TA Instruments, New Castle, DE, USA) at 298 K. Proteins and metals (CdCl_2_, NiCl_2_, MnCl_2_, CuSO_4_) were prepared in buffer A. MamM CTD was diluted to 50 μM concentration and was titrated with 1.4 mM (NiCl_2_, MnCl_2_, CuSO_4_) or 1 mM (CdCl_2_) metal solutions. The protein samples were injected into the instrument cell (170 μL) and 20 aliquots of 2.5 μL of the suitable metal solution were titrated into the cell every 350 seconds. For each protein, 3 independent titrations were measured. As a control, each metal solution was titrated into buffer A under the same experimental conditions, and all measurements were compared to a DDW-containing reference cell. Data were analysed using TA NanoAnalyze Data Analysis software, version 3.7.5 (TA Instruments, New Castle, DE, USA). The data of each measurement were fitted to an independent model combined with a blank constant model, and the given thermodynamics values are the average of the three different titrations for each protein. Errors are reported as the standard deviation.

### Site directed spin labelling

For site-directed spin labelling, protein solutions in buffer A were degassed under argon and treated with degassed 1,4-Dithiotheitol (DTT, 1 mM) for 4 hours at 4 °C with gentle agitation. Subsequently, protein samples were washed once using Zeba™ Spin desalting columns (7 kDa MWCO, 2 mL) (Thermo Fisher Scientific, Waltham, MA, USA) equilibrated with degassed buffer A. Protein was then labelled overnight at 4 °C with (1-oxyl-2,2,5,5-tetramethylpyrrolidin-3- yl)methylthiosulfonate spin label (MTSL, 20x excess) (Toronto Research Chemicals Inc., North York, Ontario, Canada) with gentle agitation. To remove excess unbound MTSL, samples were washed twice using Zeba™ Spin desalting columns equilibrated with degassed buffer A.

### Continuous wave X-band EPR spectroscopy

For metal-bound samples, freshly spin labelled C267S MamM-CTD, in buffer A at ~70 μM was used. Stock solutions of metal salts (ZnSO_4_.5H_2_O and CuCl_2_.2H_2_O) were prepared in H_2_O to a final concentration of 100 mM. Aliquots (15 μL) of spin labelled protein were incubated with a 3-fold excess of each metal salt for 40 minutes on ice. Aliquots of ~10 μL were transferred to 0.8mm (o.d.) capillary tubes for measurement.

The ambient temperature set-up for X-band cw-EPR consisted of a Bruker E500 eleXsys spectrometer fitted with an ER 4123D (dielectric RT cw-EPR) resonator. The following measuring parameters were used for data acquisition: a microwave frequency of 9.758 GHz, a modulation frequency of 100 KHz, a modulation amplitude of 1 G, and a microwave power of 0.2 mW.

### Pulsed EPR spectroscopy

Spin labelled C267S MamM-CTD samples were lyophilised and subsequently dissolved in D2O to a final concentration of 250 μM (Cd^2+^ and Fe^2+^-bound samples) or 500 μM (Mn^2+^, Ni^2+^and Cu^2+^-bound samples). All EPR samples were prepared with 30 % Glycerol-d8 (Sigma Aldrich, UK). Stock solutions of metal salts (CdCl_2_, MnCl_2_.7H_2_O, NiCl_2_, CuSO_4_.5H_2_O and Fe(NH_4_)_2_(SO_4_)_2_.6H_2_O) were prepared to a final concentration of 100 mM in D_2_O. Spin-labelled samples were incubated with a 5-fold (Cd^2+^, Mn^2+^, Ni^2+^ and Fe^2+^-bound samples) or 3-fold (Cu^2+^-bound) excess according to the dimeric protein concentration. Incubation was carried out at room temperature for 1 hour, before the addition of Glycerol-d8 to a final concentration of approximately 30 %. 80-100 μL samples were transferred to EPR quartz tubes (Wilmad SQ-707) (Wilmad-LabGlass, Vinland, NJ, USA) and flash frozen in liquid nitrogen. For the Fe^2^-bound sample, each step was carried out under a steady stream of argon to eliminate oxygen during sample preparation. X-band pulsed EPR spectra were recorded on a Bruker E580 spectrometer (Bruker, Rheinstetten, Germany) using a Bruker MD5-W1 EPR probehead equipped with a self-modified cryogen-free cryostat (Advanced Research Systems Inc., Macungie, PA, USA). The microwave pulses were amplified using a 1kW-TWT (Applied Systems Engineering Inc., Fort Worth, TX, USA). All EPR experiments were carried out at 50K. The field-swept spectrum was obtained by integrating the Hahn echo signal as a function of the magnetic field after a two-pulse sequence (10ns - τ(120ns) - 20 ns). For pulsed electron-electron double resonance (PELDOR) experiments a 4-pulse sequence was applied. Observer pulses (π/2 and π) were set to 20 ns in length, with a pump pulse of 16 ns. The frequency offset ∆ν (ν_obs_ – ν_pump_) was set to between 70-80 MHz for SL-SL measurements. The observer pulse separation τ_1_ was 120 ns in all cases apart from with Ni^2+^, where τ_1_ was 350 ns, whilst τ_2_ varied between 1300-2200 ns, and echo signals were integrated using a video amplifier bandwidth of 20 MHz. The pump pulse was stepped out in 12 ns intervals for a total number of points in T that depended upon the τ_1_ and τ_2_ used. Nuclear modulation effects were suppressed using a systematic variation of the inter-pulse delay time τ_1_ and an appropriate phase cycling routine.

PELDOR data processing was performed using both Thikhonov regularisation and DEERNet (Worswick et al., 2018) within the Matlab-based DEER Analysis (Jeschke et al., 2006) toolbox. The recently developed DEERNet method uses artificial neural networks for prediction of an inter-spin label distance distribution, and additionally gives a measure of uncertainty of the resulting distance distributions represented by 95% confidence bounds which are indicated as the grey bands in Figure 6.

