## Supplementary material for "The cation diffusion facilitator protein MamM’s cytoplasmic domain exhibits metal-type dependent binding modes and discriminates against Mn^2+^": Tables S1-S3

**<u>Supplemental information for:</u>**

[5] Lead contact

* Correspondence:

Table S1: Crystallization of MamM CTD with different metals. Related to Figure 1.

| Protein name | **MamM CTD $Cu^{2+}$-bound** | **MamM CTD $Cd^{2+}$-bound** | **MamM CTD $Ni^{2+}$-bound** |
|---|---|---|---|
| PDB code | 6GP6 | 6GMT | 6GMV |
| Crystallization conditions | 0.2 M NaCl, 0.1 M TRIS pH 8.7, 25% PEG 3350 | 0.2 M $Li_2SO_4$, 0.1 M BIS-TRIS pH 5.7, 25% PEG 3350 | 0.2 M $(NH_4)_2SO_4$, 0.1 M BIS-TRIS pH 5.5, 25% PEG 3350 |
| Cryo protectant | 50% PEG 3350 | - | - |
| Protein concentration (mg $mL^{-1}$) | 10 | 10 | 10 |
| Crystallization type | Vapor diffusion (sitting drop) | | |
| Data collection | ESRF – ID30A3 | ESRF – ID30A3 | ESRF – ID23-1 |
| Detector | Eiger X 4M | Eiger X 4M | Pilatus 6M |

Table S2: Data collection and refinement statistics of MamM CTD with different metals. Related to Figure 1.

| Protein name | **MamM CTD $Cu^{2+}$-bound [a]** | **MamM CTD $Cd^{2+}$-bound** | **MamM CTD $Ni^{2+}$-bound** |
|---|---|---|---|
| PDB code | 6GP6 | 6GMT | 6GMV |
| Data collection | ESRF – ID30A3 | ESRF – ID30A3 | ESRF – ID23-1 |
| Space group | P 2 $2_1$ $2_1$ | C 2 2 $2_1$ | C 2 2 $2_1$ |
| ***Cell dimensions*** | | | |
| a, b, c (Å) | 28.93, 73.75, 89.41 | 36.53, 94.25, 53.32 | 37.34, 94.48, 53.69 |
| α, β, γ (°) | 90, 90, 90 | 90, 90, 90 | 90, 90, 90 |
| Resolution (Å) | 2.14-44.71 (2.14-2.20) [b] | 1.59-47.12 (1.59-1.61) | 1.59-47.24 (1.59-1.62) |
| Rsym or Rmerge | 0.106 (1.863) | 0.027 (1.046) | 0.059 (2.376) |
| I/σI | 8.5 (1.0) | 20.6 (1.2) | 24.3 (1.4) |
| $CC_{1/2}$ | 0.996 (0.436) | 1.000 (0.445) | 1.000 (0.769) |
| Completeness (%) | 99.6 (96.9) | 98.2 (97.9) | 99.9 (97.4) |
| Redundancy | 5.8 (5.8) | 3.9 (4.0) | 19.4 (19.1) |
| Wavelength (Å) | 0.96771 | 0.96771 | 0.977999 |
| No. unique reflections | 11127 (864) | 12566 (597) | 13052 (616) |
| ***Refinement*** | | | |
| Resolution (Å) | 2.15-44.71 (2.15-2.46) | 1.59-47.12 (1.59-1.63) | 1.59-47.24 (1.59-1.64) |
| Rwork/Rfree | 22.01/27.51 (25.6/34.5) | 18.68/23.93 (40.0/39.9) | 18.82/22.18 (33.9/38.5) |
| *No. atoms* | | | |
| Protein | A-796, B-714 | 637 | 664 |
| Ligand/ion | 7 | 16 | 15 |
| Water | 21 | 53 | 64 |
| *B-factors* | | | |

| | | | |
|---|---|---|---|
| Protein | A-40.12, B-47.18 | 34.95 | 35.94 |
| Ligand/ion | $Cu^{2+}$-76.95, βME-50.81 | $Cd^{2+}$-43.55, $SO_4$-38.82, βME-60.39 | $Ni^{2+}$-68.24, $SO_4$-38.59, βME-73.73 |
| Water | 41.44 | 42.77 | 44.10 |
| *RMSD* | | | |
| Bond lengths (Å) | 0.009 | 0.028 | 0.031 |
| Bond angles (°) | 1.076 | 2.620 | 2.595 |
| Ramachandran statistics [c] | P: 181 (97.31%), A: 2 (1.08%), O: 3 (1.61%) | P: 76 (98.70%), A: 1 (1.30%), O: 0 (0%) | P: 71 (98.61%), A: 1 (1.39%), O: 0 (0%) |
| Missing residues | A: 211, 315-318 B: 302-318 | 211-212, 293-318 | 211, 293-318 |

Values in parentheses are for the highest resolution shell.

One crystal was used per dataset.

Datasets were collected at 100K.

[a] Collection statistics are given after the Aimless scaling; data were further processed with UCLA-DOE-LAB Diffraction Anisotropy Server(http://services.int.mbi.ucla.edu/anisoscale/).

[b] Best resolution is given for b axis (best resolution is 2.3 Å and 2.6 Å for a and c axes, respectively, after anisotropic data reduction).

[c] P- Preferred region, A- Allowed region, and O- outliers.

Table S3: Crystallographic software used for structure solution of MamM CTD with different metals. Related to Figure 1.

| Protein name | **MamM CTD $Cu^{2+}$-bound [a]** | **MamM CTD $Cd^{2+}$-bound** | **MamM CTD $Ni^{2+}$-bound** |
|---|---|---|---|
| PDB code | 6GP6 | 6GMT | 6GMV |
| Data reduction | XDS (Kabsch, 2010) | | |
| Data scaling | Aimless (Evans and Murshudov, 2013) [a] | Aimless | Aimless |
| Structure solution method | Molecular replacement – using MamM CTD WT structure (PDB code: 3W5X) | | |
| Phasing | Phaser MR (McCoy et al., 2007) | | |
| Refinement | Phenix (Adams et al., 2010) | Refmac5 (Murshudov et al., 2011) | Refmac5 |

Manual refinement was performed using Coot version 0.8.9 (Emsley et al., 2010).

Aimless, Phaser MR and Refmac5 were used through the CCP4i package (Winn et al., 2011).

[a] After Aimless scaling, data were further processed with UCLA-DOE-LAB Diffraction Anisotropy Server.
